# Identification of *bla*_GIM-1_ in *Acinetobacter variabilis* isolated from the hospital environment in Tamil Nadu, India

**DOI:** 10.1101/586164

**Authors:** Prasanth Manohar, Murugavel Ragavi, Ashby Augustine, Hrishikesh MV, Nachimuthu Ramesh

**Affiliations:** Antibiotic Resistance and Phage Therapy Laboratory, School of Bioscience and Technology, Vellore Institute of Technology (VIT), Vellore, Tamil Nadu, India.

**Keywords:** *Acinetobacter* species, *bla*_GIM-1_, carbapenem resistance, Gram-negative bacteria, German imipenemase

## Abstract

**Background:** Emergence of carbapenem resistance mechanisms among Gram-negative bacteria is a worrisome health problem. Here, we focused on to identify the presence of carbapenem-resistant bacteria among the samples collected from hospital environments in Tamil Nadu.

**Methods:** A total of 30 hospital environmental samples were collected between August 2017 and January 2018 from hospitals located in Chennai and Vellore such as lift switches, stair rails, switchboards, nursing desks, used nursing gloves, door handles, wheelchairs, touch screens, chairs and from pillars inside the hospitals.

**Results and discussion:** A total of 22 carbapenem-resistant Gram-negative bacteria were isolated that included *Escherichia coli, Klebsiella* sp., *Enterobacter* sp., *Salmonella* sp., *Pseudomonas aeruginosa* and *Acinetobacter* sp. Interestingly, *bla*_GIM-1_ was detected in *Acinetobacter variabilis* strain isolated in samples collected from hospitals. Unlike other studies, the identified GIM-1 was not plasmid encoded, and this is the first report for the presence of GIM-1 (German imipenemase) in India.

**Conclusion:** Extensive surveillance programs are necessary to trace the uncontrolled spread of carbapenem-resistance genes in order to reduce the rapid spread of resistance.

## Introduction

Antibiotic resistance has become one of the major clinical and public health concerns [1]. The spread of antibiotic resistance in clinical settings (nosocomial infections) is a well-recognized problem, and this becomes overlooked with great importance [1]. Multidrug-resistant bacteria (MDR) are spreading at an uncontrolled rate, especially in hospital areas [2]. In recent years, it is found that there is a prevalence of a large population of multi-drug resistant bacteria in and around the hospital environment [1]. These are bacteria which are exposed to a variety of antibiotics and simultaneously becoming resistant, which makes a major threat to the public health [3]. Carbapenem is a subgroup in beta-lactam antibiotics which is considered as one of the last hope for the threatening multi-drug resistant bacterial infections [4,5]. But developing carbapenem-resistant Gram-negative bacteria is one of the serious concerns. More dangerously, an MDR bacterium having one or more beta-lactamase genes can involve in horizontal transfer of resistance genes to other bacterial species or strains [6]. Some of the prevalent carbapenem-resistance genes in India are; New Delhi Metallo-beta-lactamase (NDM), Imipenemase (IMP), Verona Integron-Mediated metallo-beta-lactamase (VIM), Oxacillinase (OXA) [7]. In another study, the identification of DIM-1 gene in *Pseudomonas aeruginosa* was reported for the first time in India by our group [8]. Antibiotic resistance surveillance programs in hospitals are essential to monitor and control the spread of antibiotic-resistant infections (nosocomial). Community-associated or community onset of carbapenem-resistant bacterial infections is reported to have a prevalence of 0-30% [9]. The main focus of this study is to identify the presence of carbapenem-resistant bacteria from samples collected in the hospital environment.

## Methods

In the present study, a total of 30 samples were collected from in and around hospitals in Chennai and Vellore, Tamil Nadu during the period between August 2017 and August 2018. The samples were taken from lift switches, stair rails, switchboards, nursing desks, used nursing gloves, door handles, wheelchairs, touch screens, chairs and pillars inside the hospital. Sterile swabs dipped in sterile tryptone broth were used for sample collection, and collected samples were immediately transported and processed at Antibiotic Resistance and Phage Therapy Laboratory, VIT, Vellore. Bacterial isolation was carried out using the spread plate technique using MacConkey agar M081 (Himedia, India) amended with 8 µg/mL of meropenem. Minimal inhibitory concentration (MIC) was performed using micro-broth dilution method as per the CLSI guidelines. Modified Hodge test (MHT) was performed to evaluate the carbapenemase production as explained elsewhere [7]. Bacterial identification was carried out with the VITEK identification system (bioMerieux) and by 16S rRNA analysis using universal primers 27F and 1492R. DNA was isolated using a boiling lysis method as explained elsewhere [7]. Further, molecular screening of beta-lactamase genes such as *bla*_VIM_, *bla*_IMP_, *bla*_KPC_, *bla*_NDM_, *bla*_OXA-48_, *bla*_OXA-1_, *bla*_OXA-4_, *bla*_OXA-30_, *bla*_OXA-23_, *bla*_OXA-24,_ *bla*_OXA-51_, *bla*_OXA-58_, *bla*_AIM_, *bla*_GIM_, *bla*_DIM_, *bla*_SIM_, *bla*_BIC_ and *bla*_GES_ were performed using multiplex PCR using the primers and PCR conditions as explained earlier [7, 8]. Plasmid DNA was isolated (only for the isolates that were identified to harbour resistance genes) using HiPurA Plasmid DNA Miniprep Purification kit (Himedia, India). All the PCR products were sequenced (Eurofins Genomics India Pvt. Ltd., Bangalore) and sequence results were analysed against available BLASTN sequences and deposited in Genbank.

## Results

A total of 22 non-repetitive, carbapenem-resistant, Gram-negative bacteria were isolated from 30 samples collected from hospital environments. Of the 22 carbapenem-resistant isolates, 15 were identified as lactose fermenters, and 7 were non-fermenters. The isolates were identified as *E. coli* (n=7), *Klebsiella* species (n=4), *Enterobacter* species (n=4), *Salmonella* species (n=2), *Acinetobacter* species (n=2) and *Pseudomonas aeruginosa* (n=3). The MIC results for meropenem/ imipenem were ranged between 4 µg/mL and >128 µg/mL (Table 1). None of the isolates was found to be positive for carbapenemase production by MHT. Molecular characterization results showed the presence of a *bla*_GIM_ gene in *Acinetobacter variabilis*. Genes *bla*_VIM_, *bla*_IMP_, *bla*_KPC_, *bla*_NDM_, *bla*_OXA-48_, *bla*_OXA-1_, *bla*_OXA-4_, *bla*_OXA-30_, *bla*_OXA-23_, *bla*_OXA-24,_ *bla*_OXA-51_, *bla*_OXA-58_, *bla*_AIM_, *bla*_DIM_, *bla*_SIM_, *bla*_BIC_ and *bla*_GES_ were absent in all the isolates tested. Plasmid DNA was isolated from *Acinetobacter variabilis* and presence of ≈20kb plasmid was noted. Further screening for the presence of *bla*_GIM_ gene using plasmid DNA confirmed that *bla*_GIM_ gene was bound to chromosomal DNA, not in plasmid DNA. Sequencing results confirmed that the amplified *bla*_GIM_ gene is *bla*_GIM-1_, and to the best of our knowledge this is a first report for the presence of *bla*_GIM-1_ in India. [Accession number: MG764090]

**Table 1:**
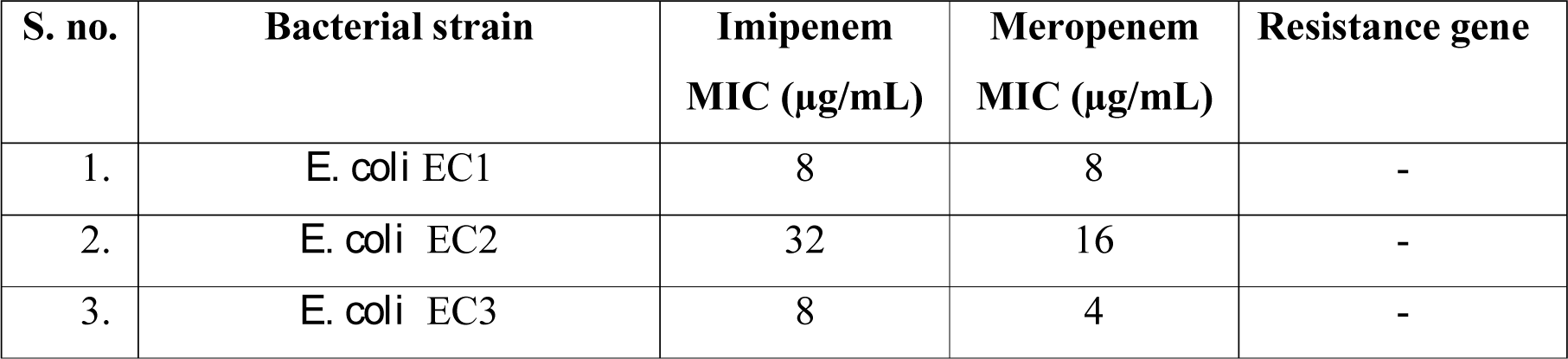
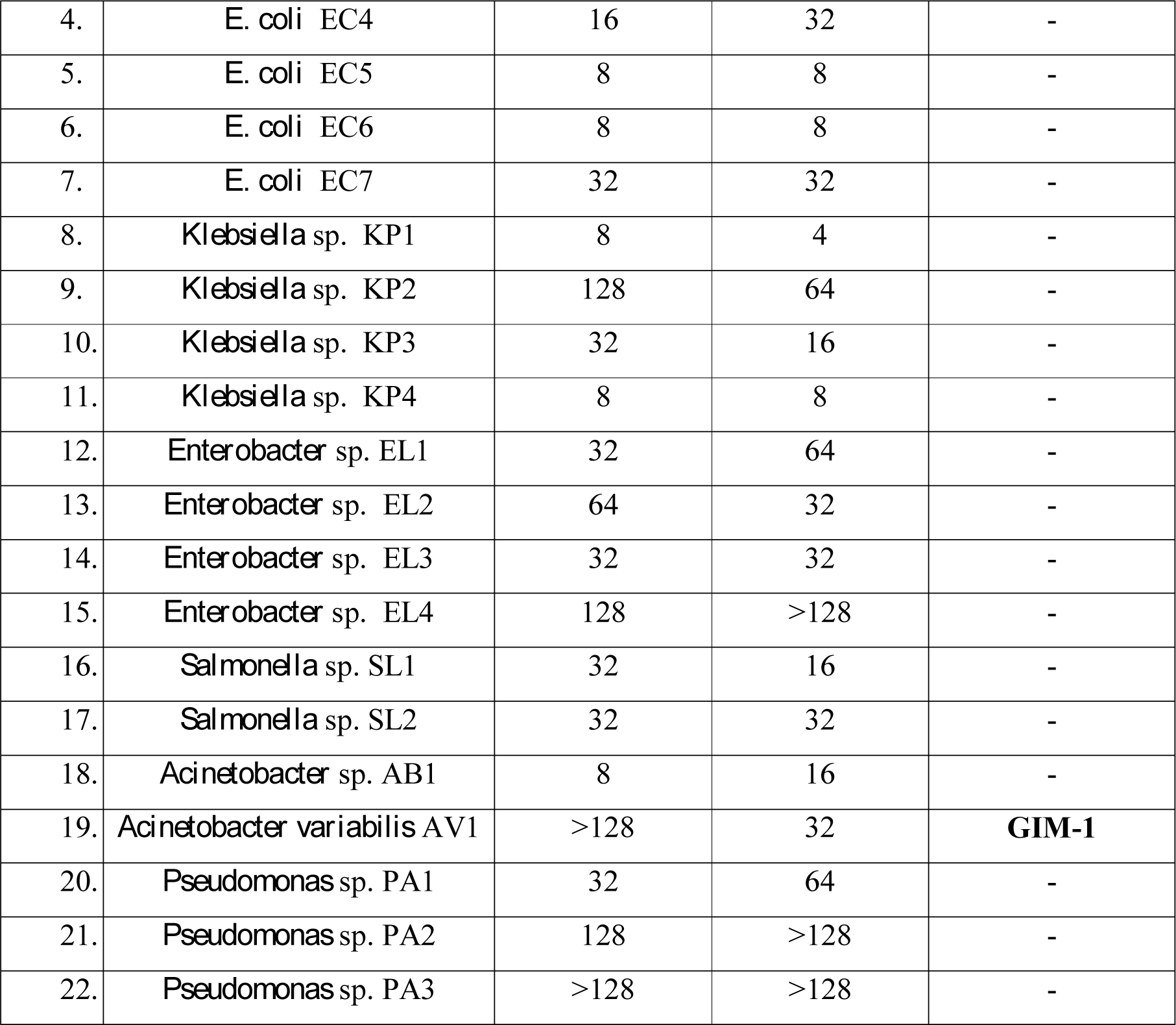
Classification of resistance among the Gram-negative bacteria isolated from hospital environmental samples.

## Discussion

Carbapenem-resistance among Gram-negative bacteria is one of the increasing public health concerns. Nosocomial infections are considered as one of the serious health care problems, notably; the rapid spread of antibiotic-resistant bacterial infections is scary [10]. In India, the prevalence of carbapenem-resistant bacteria is being reported and multi-drug resistant bacterial infections are on the rise [7]. Importantly, this study reports the emergence of carbapenem-resistant bacteria in the hospital environment and presence of GIM-1 gene in *A. variabilis.* In the past, GIM (German Imipenemase)-type carbapenem-resistance gene was reported in *P. aeruginosa* in 2002 and in common, the reports of GIM genes were from the bacteria isolated in and around Germany. To the best of our knowledge, this is the first study to report the emergence of GIM-1 from India or even from the Asian subcontinent. Earlier, the presence of GIM-1 genes was reported to be encoded on plasmids, and also in mobile genetic elements such as integrons [11]. This is one of the rarest studies to report the presence of GIM-1 gene in *A. variabilis* isolated from environmental samples, and the resistance gene was bound to chromosomal DNA and not in a plasmid. It is very common to identify GIM-1-type carbapenem resistance mechanism in *Enterobacteriaceae* and *P. aeruginosa*, but the prevalence of GIM-type carbapenemases in *Acinetobacter* is very rare. The isolation of *A. variabilis* in India indicates that GIM-1 is no longer confined to one geographical region as previously thought and these are more widespread among other non-*Enterobacteriaceae*. This clearly indicates the potential transmission of GIM-1-type carbapenemase genes among other Gram-negative bacteria. The increasing prevalence of carbapenem-resistant Gram-negative bacteria (CR-GNB) in the hospital environment, causing nosocomial infections, is challenging particularly during the clinical recovery of patients. This study opens the door for the detection of new carbapenemase genes in India; therefore, extensive resistance surveillance is needed to learn more about the emergence of GIM-1-producing bacteria in India.

## Ethical Statement

The ethical approval for this study was obtained from Institutional Biosafety Committee “Ref. No. VIT/IBSC/07/October, 2018”.

## Author’s contribution

PM designed the study and PM, MR, AA, HMV executed the study. PM analysed the data and validated the data. PM and MR wrote the manuscript draft. NR confirmed the research data and NR finalized the manuscript.

